# theseus - An R package for the analysis and visualization of microbial community data

**DOI:** 10.1101/295675

**Authors:** Jacob R. Price, Stephen Woloszynek, Gail Rosen, Christopher M. Sales

## Abstract

**theseus** is a collection of functions within the R programming framework [1] to assist microbiologists and molecular biologists in the interpretation of microbial community composition data.

## 1 Introduction

Ecological analysis of community composition data requires a variety of tools and approaches including, but not limited to, diversity summary statistics, regression modeling, constrained and unconstrained ordination, clustering, differential abundance detection and testing, and community network analysis. In addition, researchers must often make decisions about preprocessing, filtering, and denoising their data, prior to carrying out analysis. The use of amplicon sequencing data in the exploration of microbial communities has dramatically increased as a result of improvements to, and reduced costs of, high-throughput sequencing technologies.

A number of open-source packages available in the R programming framework [1], such as the **vegan** [2], **phyloseq** [3], **microbiome** [4], and **themetagenomics** [5] R packages, have been developed to address not only traditional community datasets, but also the unique challenges associated with amplicon sequencing data. Here we introduce a new R package titled **theseus** which builds upon the functionality that is offered by the proceeding community analysis packages and expands upon their methods by offering new ways to create, analyze, and interpret raw data and results.

## 2 Functional Description and Advantages

### 2.1 Included Data

Beyond providing R functionality, the authors intend to use the **theseus** package as a vehicle for disseminating research products (such as **phyloseq** objects and other datasets) for a variety of uses, including demonstrating package functionality, instruction in academic courses, and, especially, encouraging replication of experimental results and further exploration of those results. The authors expect to continue to expand the available datasets within **theseus** as research products are published and functionality is expanded.

**theseus** is currently (version 0.1.0) packaged with datasets generated by Price et. al. (2018) [6] while investigating the impact of wastewater treatment plant (WWTP) effluent on receiving streams. The principle dataset from this study is a **phyloseq** object (WWTP Impact) that contains 16S rDNA amplicon sequencing data from samples collected at 6 sites on 2 days (12 total samples) along Wissahickon Creek and Sandy Run in southeastern Pennsylvania, USA. Sequencing data was processed as described in Price et. al. (2018) [6], and combined with chemical data to create a **phyloseq** object. The raw data and scripts used to generate this **phyloseq** object (as well as the **phyloseq** object itself) can be obtained from the authors’s GitHub repository (https://github.com/JacobRPrice/WWTP_Impact_on_Stream). It is this dataset that will be used to present functionality from the **theseus** package below.

### 2.2 qualcontour

Trimming and quality filtering is one of the first steps taken when analyzing amplicon sequencing results. Indiscriminate trimming can cause a host of issues for downstream processes. Function qualcontour is intended to assist the user with deciding where trimming should be performed.

qualcontour’s (quality contour) two required arguments are character vectors of the file paths for forward (*f path*) and reverse (*r path*) reads. qualcontour tabulates the distribution of quality scores at each read cycle for the forward and reverse reads independently and then averages (arithmetic mean) the quality scores for each (forward/reverse) cycle combination. These values are then plotted as a **ggplot2** object.

Using this visualization (Figure 1), we can see that reverse read quality drops earlier, and more gradually when compared with the forward read quality. Selecting the appropriate trimming location can be tricky and there are multiple ways to approach the challenge. Two places (or rules) to start with are:

**Figure 1.**
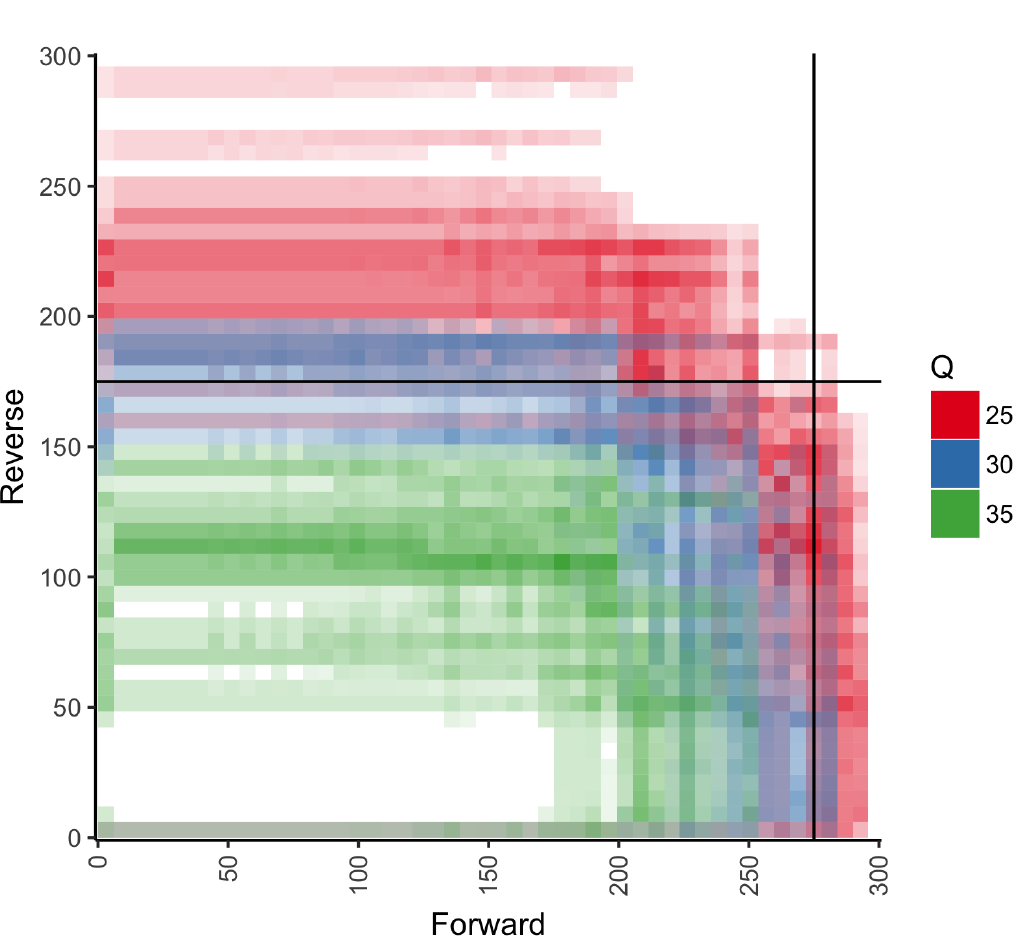
Function qualcontour is intended to help researchers select trimming locations based upon quality profiles of both forward and reverse reads.

- trimming where a severe/sudden drop in read quality occurs, such as at 275 for the forward read
- trimming where the quality scores drop below a certain (sometimes arbitrarily selected) value, such as 175 in the reverse reads, where we estimate that read quality hit *Q* = 30.

### 2.3 prev

Low count taxa are often filtered from OTU tables before analysis to reduce (possible) error or noise. Examination of the raw (unfiltered) OTU table should be carried out to ensure that appropriate thresholds for prevalence (number of samples a taxa was observed in) and abundance (the total number of times a taxa was observed) are being selected. Function prev (prevalence), adapted from [7], plots each taxon according to their prevalence and abundance within the dataset (Figure 2). Users can designate values for minimum abundance (total taxa counts in the dataset) and prevalence (number of samples in which the taxa was observed).

**Figure 2.**
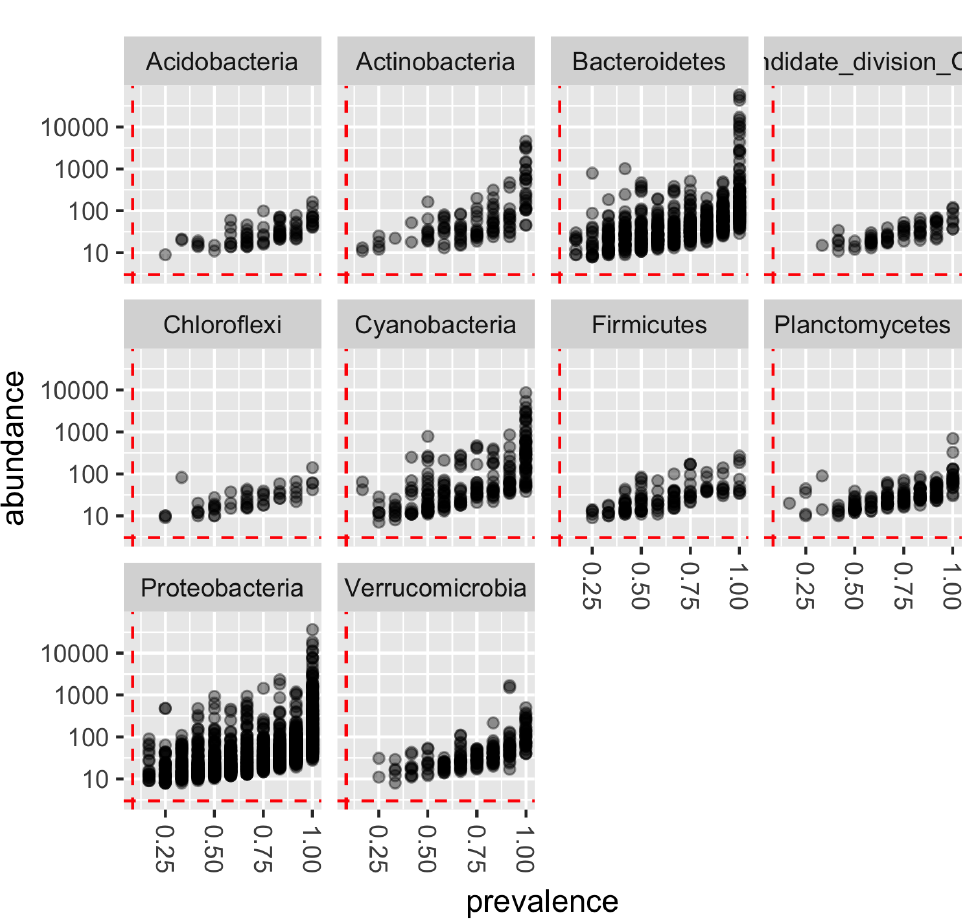
Function prev can be used to identify filtering/denoising parameter values.

### 2.4 envtoverlay

The calculations for unconstrained ordination do not take environmental or sample metadata into account. The **vegan** functions envfit and ordisurf attempt to fit environmental descriptors into ordination space and display them as vectors and surfaces respectively. Gavin Simpson, one of **vegan**’s authors, has a great blog entry explaining how these functions work and are interpreted. Jari Oksanen has provided several vignettes on **vegan**’s homepage that also discuss the application of these functions. Function envtoverlay (environmental variable overlay) takes advantage of **ggplot2** to make these plots (usually generated in base::plot) more aesthetically pleasing and user friendly, particularly when the user would like to present the results as a faceted plot (Figure 3). The user provides a character vector of covariates upon which the envfit and ordisurf functions to be applied.

**Figure 3.**
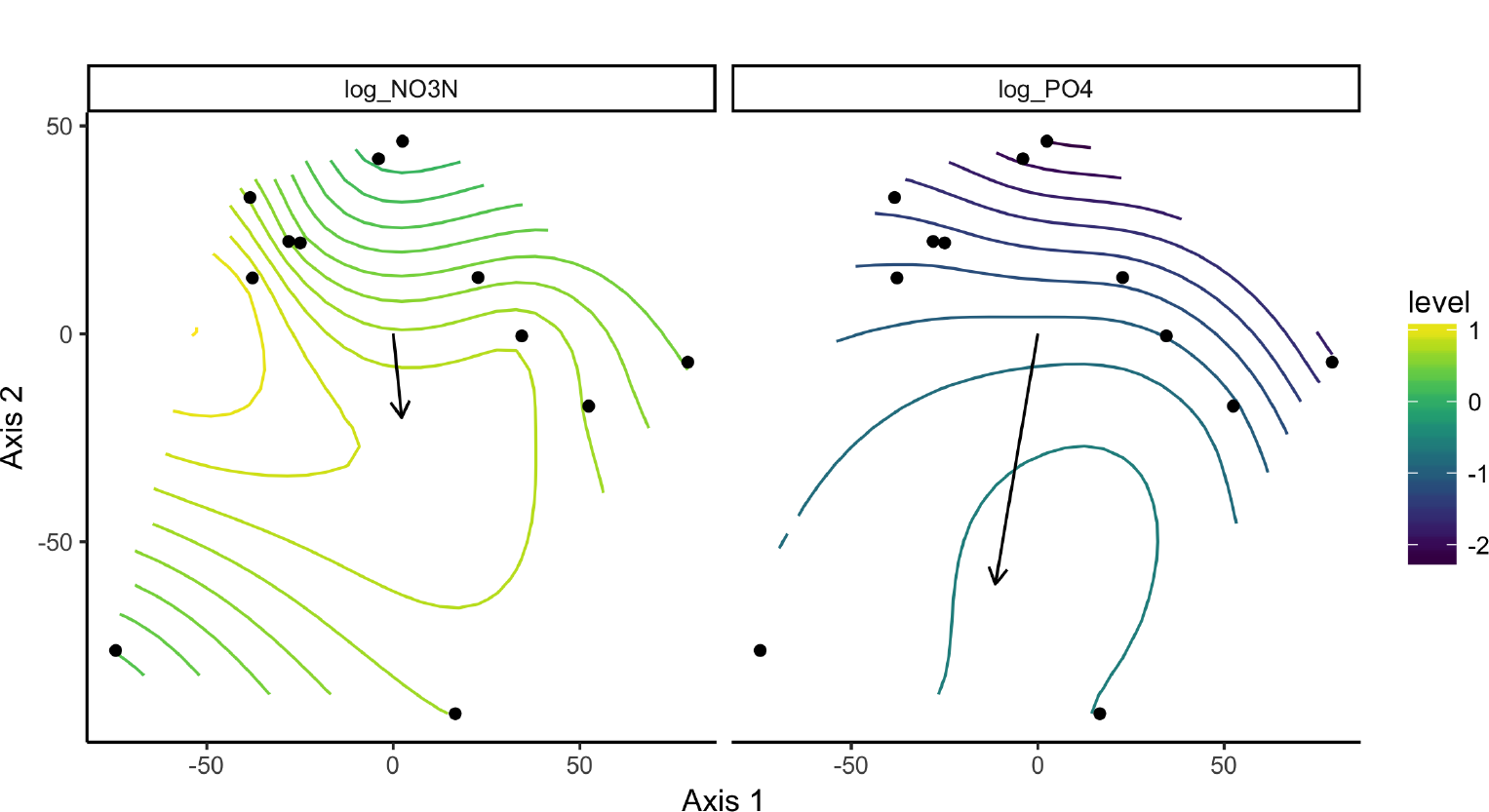
Function envtoverlay generates attractive figures using **vegan**’s envfit and ordisurf functions.

### 2.5 constord

Function constord (constrained ordination) carries out constrained ordination on a **phyloseq** object and plots the results. Constrained correspondence analysis (‘CCA’) and redundancy analysis (‘RDA’) are the two methods currently implemented within this function. Figure 4 presents a RDA analysis of the WWTP_Impact dataset using log-scaled nitrate (log_NO3N) and phosphate (log_PO4) as constraining variables; scale=2 was selected in the generation of this figure.

**Figure 4.**
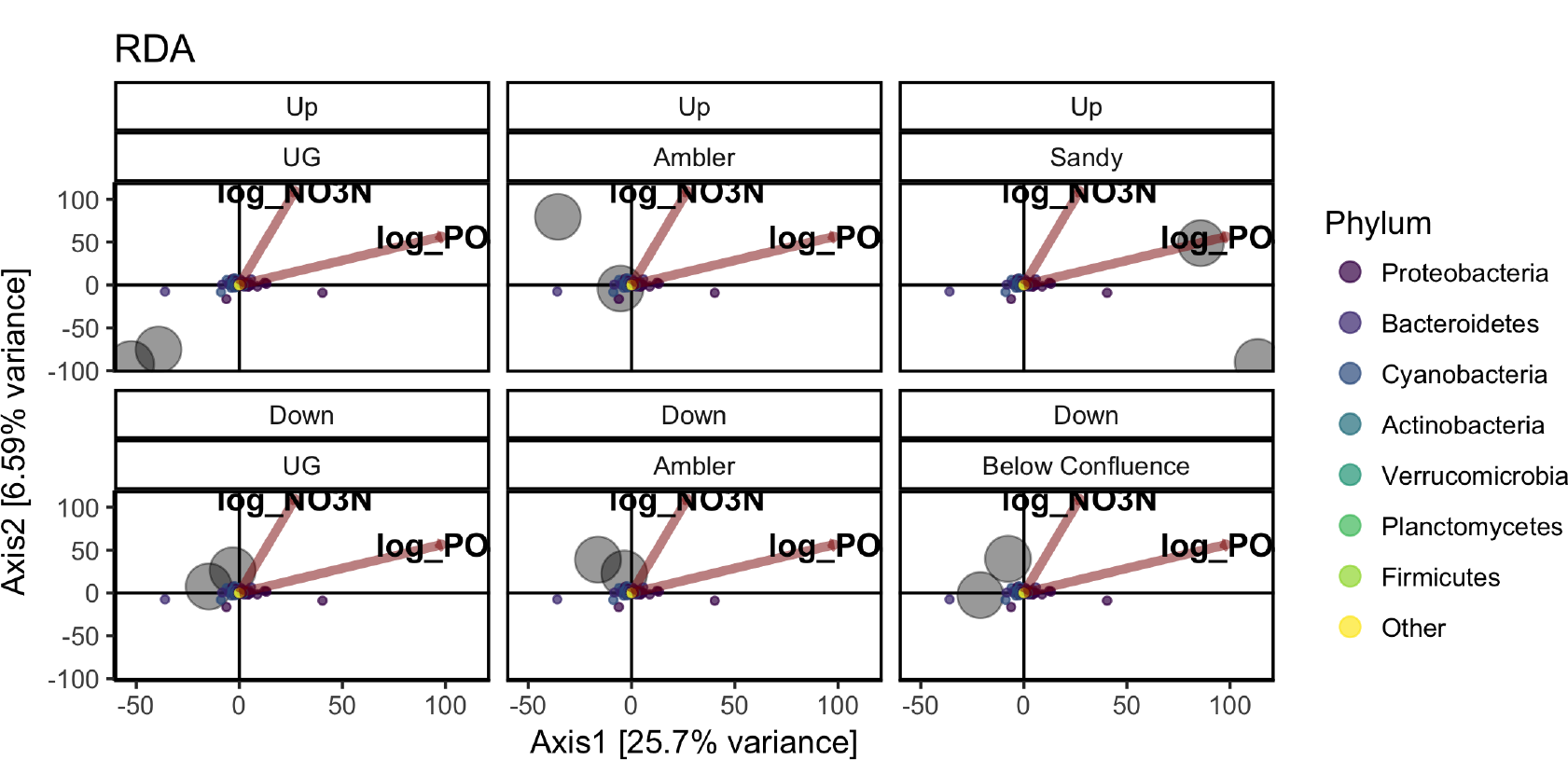
Function constord carries out constrained ordination (CCA or RDA) and plots the results.

There are several differences between constord and phyloseq::plot ordination, and they each have their own strengths. The highlights of constord are:

- constraining variables are included in the constord plot, a feature not currently present in **phyloseq**’s approach, but still possible with some extra coding.
- constord has an argument *scaling* which allows the user to select whether species scaling (1) or site scaling (2) is used when returning the scores to be plotting. Currently, phyloseq::plot ordination returns site scaling (2). The choice of scaling is important and should be selected depending upon whether the goal is to compare the arrangement of sites or species.
- The aspect ratio of the ordination plots themselves are scaled according to the ordination’s eigenvalues to more accurately represent the distances between sites/samples/taxa, as described by Callahan et. al. (2016) [7].

**Note:**

- The numbering approach is consistent with **vegan** (**theseus** and **phyloseq** rely on vegan::scores to get the scores themselves).
- The options for this function and its plot are evolving and we may include additional ordination methods and plotting features later on.

### 2.6 cohort relabund

A number of methods exist to analyze and identify differential gene expression datasets; some of them are also appropriate for analyzing microbiome data. The **DESeq2** R package [8] is one such analytical method. The results from **DESeq2** enable researchers to test, compare, and contrast microbial abundances using models/designs as simple or complex as the experiment allows. One difficultly with working with these results is interpreting the results of multiple comparisons/tests simultaneously, i.e., integrating two or more comparisons in a holistic view of how microbiomes are changing. We have created a graphical representation which we call a cohort relative abundance plot (hence the function name cohort relabund) to assist in the interpretation of multiple comparisons.

For each comparison, an OTU can can be categorized into one of 3 categories: (statistically significant) decreased, no (statistically significant) change, and (statistically significant) increased. If a researcher wishes to make 2 *simultaneous* comparisons, each taxon could fall into one of 9 possible categories (what we call cohorts): decreased (comparison 1) / decreased (comparsion 2), decreased (comparison 1) / no change (comparsion 2), decreased (comparison 1) / increased (comparsion 2), etc. Function cohort_relabund allows users to provide a **phyloseq** object and two comparison objects to generate a 3x3 faceted plot showing how each taxon is partitioned between the 9 possible cohorts Figure 5.

**Figure 5.**
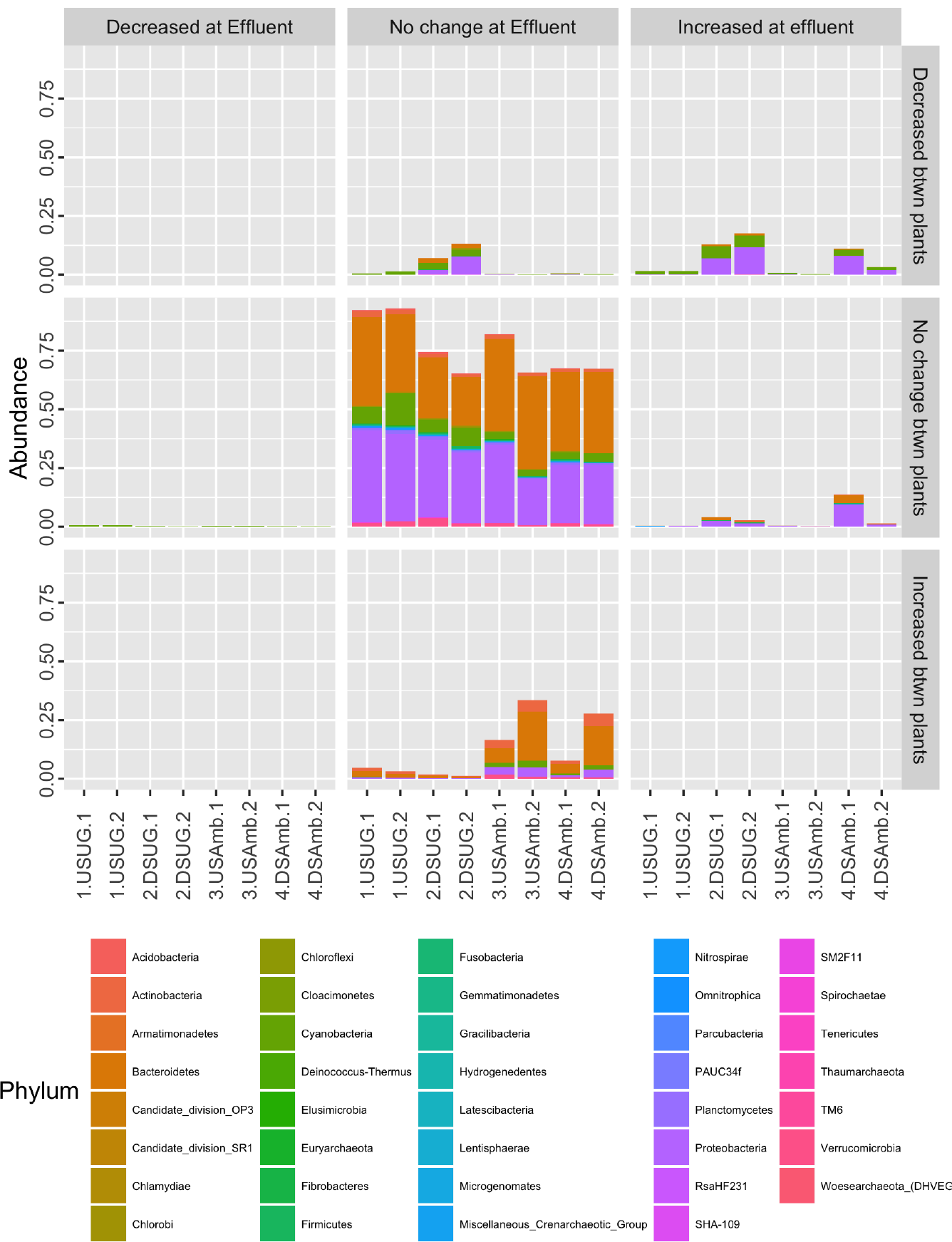
Function cohort relabund enables researchers to easily interpret OTU abundances over two contrasting comparisons.

## 3 Software Availability

Version 0.9.0 of the **theseus** R package is available through the Comprehensive R Archive Network (CRAN) or from Drexel’s Ecological & Evolutionary Signal-Processing & Informatics (EESI) Lab’s GitHub repository (https://github.com/EESI/theseus).

## Acknowledgements

Much of **theseus**’ functionality relies on functions and approaches provided by **phyloseq** [3] and **vegan** [2].

